# Optimal Design of Single-Cell Experiments within Temporally Fluctuating Environments

**DOI:** 10.1101/812479

**Authors:** Zachary R Fox, Gregor Neuert, Brian Munsky

**Affiliations:** Inria Saclay Ile-de-France, Palaiseau 91120, France; Institut Pasteur, USR 3756 IP CNRS Paris, 75015, France; School of Biomedical Engineering, Colorado State University Fort Collins, CO 80523, USA; Department of Molecular Physiology and Biophysics, School of Medicine, Vanderbilt University, Nashville, TN 37232, USA; Department of Biomedical Engineering, School of Engineering, Vanderbilt University, Nashville, TN 37232, USA; Department of Pharmacology, School of Medicine, Vanderbilt University, Nashville, TN 37232, USA; Department of Chemical and Biological Engineering, Colorado State University Fort Collins, CO 80523, USA

## Abstract

Modern biological experiments are becoming increasingly complex, and designing these experiments to yield the greatest possible quantitative insight is an open challenge. Increasingly, computational models of complex stochastic biological systems are being used to understand and predict biological behaviors or to infer biological parameters. Such quantitative analyses can also help to improve experiment designs for particular goals, such as to learn more about specific model mechanisms or to reduce prediction errors in certain situations. A classic approach to experiment design is to use the Fisher information matrix (FIM), which quantifies the expected information a particular experiment will reveal about model parameters. The Finite State Projection based FIM (FSP-FIM) was recently developed to compute the FIM for discrete stochastic gene regulatory systems, whose complex response distributions do not satisfy standard assumptions of Gaussian variations. In this work, we develop the FSP-FIM analysis for a stochastic model of stress response genes in *S. cerevisae* under time-varying MAPK induction. We verify this FSP-FIM analysis and use it to optimize the number of cells that should be quantified at particular times to learn as much as possible about the model parameters. We then extend the FSP-FIM approach to explore how different measurement times or genetic modifications help to minimize uncertainty in the sensing of extracellular environments, and we experimentally validate the FSP-FIM to rank single-cell experiments for their abilities to minimize estimation uncertainty of NaCl concentrations during yeast osmotic shock. This work demonstrates the potential of quantitative models to not only make sense of modern biological data sets, but to close the loop between quantitative modeling and experimental data collection.

## INTRODUCTION

The standard approach to design experiments has been to rely entirely on expert knowledge and intuition. However, as experimental investigations become more complex and seek to examine systems with more subtle non-linear interactions, it becomes much harder to improve experimental designs using intuition alone. This issue has become especially relevant in modern single-cell-single-molecule investigations of gene regulatory processes. Performing such powerful, yet complicated experiments involves the selection from among a large number of possible experimental designs, and it is often not clear which designs will provide the most relevant information. A systematic approach to solve this problem is model-driven experiment design, in which one combines existing knowledge or experience to form an assumed (and partially incorrect) mathematical model of the system to estimate and optimize the value of potential experimental settings. In practice, such preliminary models would be defined by existing data taken in simpler or more general settings such as inexpensive bulk experiments, or would be estimated from literature values conducted on similar genes, pathways or organisms. When parameter or model structures are uncertain these could be described according to a prior distribution, and experiments would need to be selected according to which performs best on average across the many possible model/parameter combinations.

In recent years, model-driven experiment design has gained traction for biological models of gene expression, whether in the Bayesian setting [1] or using Fisher information for deterministic models [2], and even in the stochastic, single-cell setting [3–7]. Despite the promise and active development of model-driven experiment design from the theoretical perspective, more general, yet biologically-inspired approaches are needed to make these methods suitable for the experimental community at large. In this work, we apply model-driven experiment design to an experimentally validated model of stochastic, time-varying High Osmolarity Glycerol (HOG) Mitogen Activated Protein Kinase (MAPK) induction of transcription during osmotic stress response in yeast [8–10]. To demonstrate a concrete and practical application of model-driven experiment design, we find the optimal *measurement schedule* (i.e., when measurements ought to be taken) and the appropriate *number of individual cells* to be measured at each time point.

In our computational analyses, we consider the experimental technique of single-mRNA Fluorescence *in situ* Hybridization (smFISH), where specific fluorescent oligonucleotide probes are hybridized to mRNA of interest in fixed cells [11, 12]. Cells are then imaged, and the mRNA abundance in each cell are counted, either by hand or using automated software such as [13]. Such counting can be a cumbersome process, but little thought has been given typically to how many cells should be measured and analyzed at each time. Furthermore, when a dynamic response is under investigation, the specific times at which measurements should be taken (i.e., the times after induction at which cells should be fixed and analyzed) is also unclear. In this work, we use the newly developed finite state projection based Fisher information matrix (FSP-FIM, [6]) to optimize these experimental quantities for osmotic stress response genes in yeast.

The first part of our current study introduces a discrete stochastic model to analyze time-varying MAPK-induced gene expression response in yeast and then demonstrates the use of FSP based Fisher information to optimize experiments to minimize the uncertainty in model parameters. In the second part of this study, we expand upon this result to find and experimentally verify the optimal smFISH measurement times and cell numbers to minimize uncertainty about unknown environmental inputs (e.g., salt concentrations) to which the cells are subjected. In this way, we are presenting a new methodology by which one can optimally examine behaviors of natural cells to obtain accurate estimations of environmental changes.

## BACKGROUND

Gene regulation is the process by which small molecules, chromatin regulators, and general and gene-specific transcription factors interact to regulate the transcription of DNA into RNA and the translation of mRNA into proteins. Even within populations of genetically identical cells, these single-molecule processes are stochastic and give rise to cell-to-cell variability in gene expression levels. Adequate description of such variable responses can only be achieved through the use of stochastic computational models [14–17]. In the following subsections, we first introduce a non-equilibrium discrete stochastic model of HOG1-MAPK-induced gene expression, and we then discuss how this model can be analyzed and compared to data using finite state project analyses. All analysis codes are available at https://github.com/MunskyGroup/Fox_Complexity_2020.

### Discrete stochastic model of HOG1-MAPK-induced gene expression

To motivate and demonstrate our new approach, we focus our examination on the dynamics of the HOG1-MAPK pathway in yeast, which is a model system to study osmotic stress driven dynamics of signal transduction and gene regulation in single cells [18–23]. Discrete stochastic models of HOG1-MAPK activated transcription have been used successfully to predict the variability in adaptive transcription responses across yeast cell populations [9, 10, 24]. In particular, the authors in [9] used smFISH data to fit and cross-validate a number of different potential models with different numbers of gene states and time varying parameters. They found that dynamics of two stress response genes, *STL1* and *CTT1*, could each be described accurately by the model depicted in Fig. 1a.

**FIG. 1.**
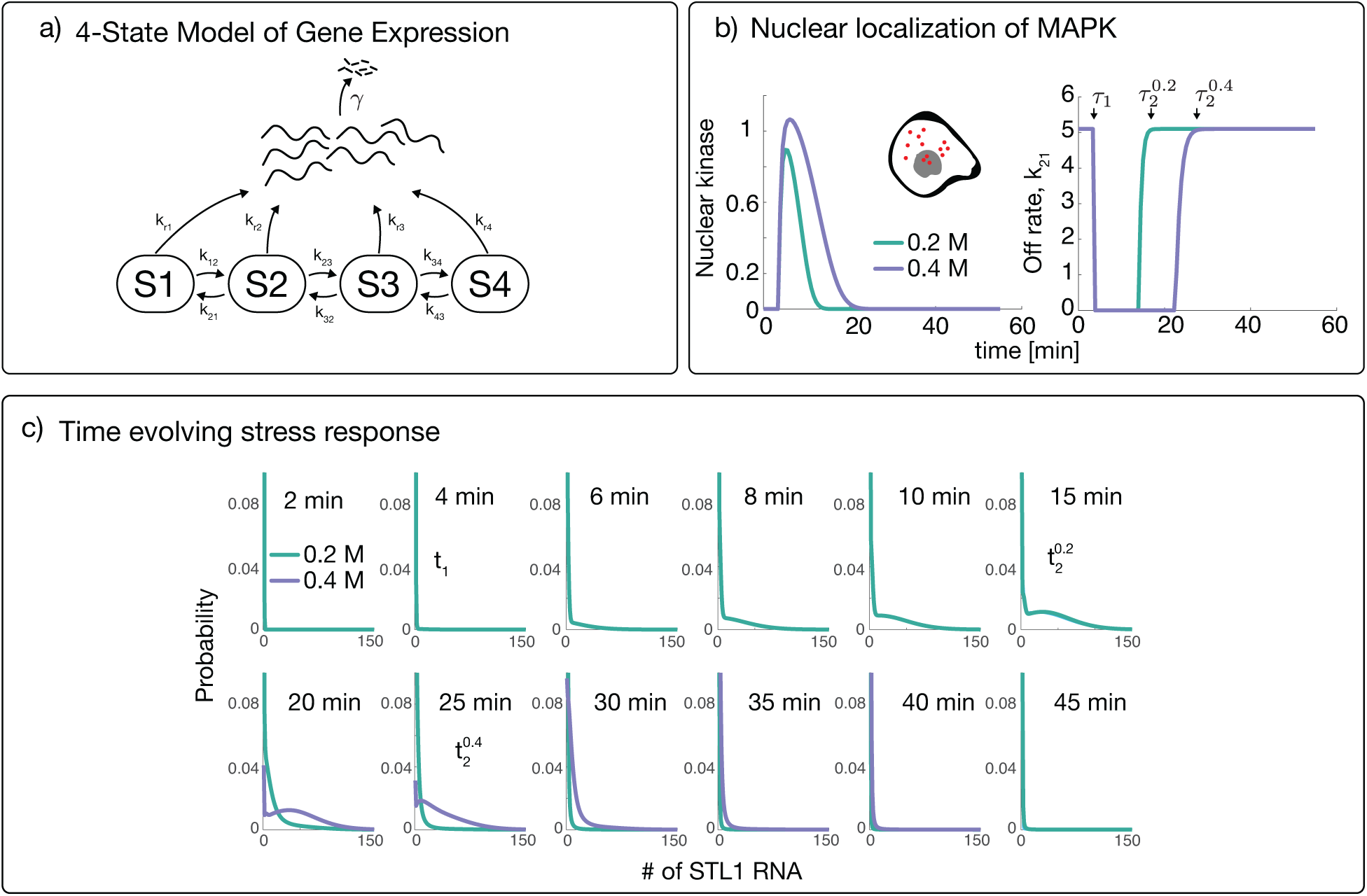
Stochastic modeling of osmotic stress response genes in yeast. (a) Four-state model of gene expression, where each state transcribes mRNA at a different transcription rate, but each mRNA degrades at a single rate *γ*. (b) The effect of measured MAPK nuclear localization (depicted as red dots in the cell) (left) on the the rate of switching from gene activation state S2 to S1 (right) under 0.2M or 0.4M NaCl osmotic stress. The time at which *k*_21_ turns off is denoted with *τ*_1_ and is independent of the NaCl level. The time at which *k*_23_ turns back on is given by 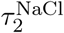 depending on the level of NaCl. (c) Time evolution of the *STL1* mRNA in response to the 0.2M and 0.4M NaCl stress. Model and parameters from [10] and summarized in Supplementary Notes I and II and Supplementary Tables I and II.

In brief, the model [9] consists of transitions between four different gene states (S1, S2, S3, and S4). The probability of a transition from the *i*^th^ to the *j*^th^ gene state in the infinitesimal time *dt* is given by the propensity function, *k*_*ij*_*dt*. Most of the rates {*k*_ij_} are constant in time, except for the transition from S2 to S1, which is controlled by the time-varying level of the HOG1-MAPK signal in the nucleus, *f* (*t*). The resulting time-varying rate *k*_21_ is defined using a linear threshold function,

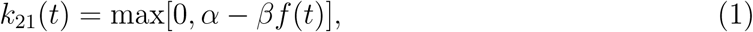

where *α* and *β* set the threshold for *k*_21_(*t*) activation/deactivation. The function *f* (*t*) was calibrated at several NaCl concentrations by fitting the HOG1-MAPK nuclear localization signals as measured using a yellow fluorescence protein reporter [10]. Figure 1b (left) shows *f* (*t*) for osmotic stress responses to 0.2M and 0.4M NaCl, and Fig. 1b (right) shows the corresponding values of *k*_21_(*t*). In addition to the state transition rates, each *i*^th^ state also has a corresponding mRNA transcription rate, *k*_ri_. All mRNA molecules degrade with rate *γ*, independent of gene state. Further descriptions and validations of this model are given in Supplementary Note 1 and in [9, 10, 24]. All experimentally determined parameters for the *STL1* and *CTT1* transcription regulation models are provided in Supplemental Table S1, and experimentally determined parameters for the HOG1-MAPK Signal Model are listed in Supplemental Table S2 [10].

### The Finite State Projection analysis of stochastic gene expression

To analyze the model described above, we apply the chemical master equation (CME) framework of stochastic chemical kinetics [25]. Combining the time-varying and constant state transition rates {*k*_*ij*_}, transcription rates {*k*_ri_}, and degradation rate *γ* from above, the CME can be written in matrix form as a linear ordinary differential equation, 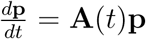, where the time-varying matrix **A**(*t*) is known as the infinitesimal generator (See Supplementary Note 1). The CME has been the workhorse of stochastic modeling of gene expression, and it is usually analyzed using simulated sample paths of its solution via the stochastic simulation algorithm [26] or with moment approximations [8, 27]. Alternatively, the CME can also be solved with guaranteed errors using the FSP approach [28, 29], which reduces the full CME only to describe the flow of probability among the most likely observable states of the system. Details of the FSP approach to solving chemical kinetic systems are provided in Supplementary Note 1. Application of the FSP analysis to the model in Fig. 1a with time varying rates *k*_21_ from Fig. 1b predicts time-evolving probability distributions as shown in Fig. 1c [10].

### Likelihood of smFISH data for FSP models

Recently, it has come to light that for some systems, it is critical to consider the full distribution of biomolecules across cellular populations when fitting CME models [6, 10]. To match CME model solutions to single-cell smFISH data, one needs to compute and maximize the likelihood of the data given the CME model [9, 10, 24, 30]. Fortunately, the FSP approach allows for computation of the likelihood with guaranteed accuracy bounds [28]. We assume that measurements at each time point 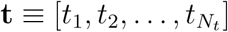 are independent, as justified by the fact that fixation of cells for measurement precludes temporal cell-to-cell correlations. Measurements of *N*_*c*_ cells can be concatenated into a matrix 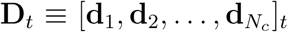 of the observable mRNA species at each measurement time *t*.

The likelihood of making the independent observations for all *N*_*c*_ measured cells is the product of the probabilities of observing each cell’s measured state. For most gene expression models, however, states are only partially observable, and we define the observed state 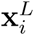 as the marginalization (or lumping) over all full states {**x**_*j*_}_*i*_ that are indistinguishable from **x**_*i*_ based on the observation. For example, the model of *STL1* transcription consists of four gene states (S1-S4, shown in Fig. 1a), which are unobserved, and the measured number of mRNA, which is observed. If we let index *i* denote the number of mRNA, then the observed state 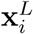 would lump together the full states (S1,*i*), (S2,*i*), (S3,*i*), and (S4,*i*). We next define *y*_*i*_ as the number of experimental cells that match 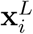 at time *t*. Under these definitions, the likelihood of the observed data (and its logarithm) given the model can be written:

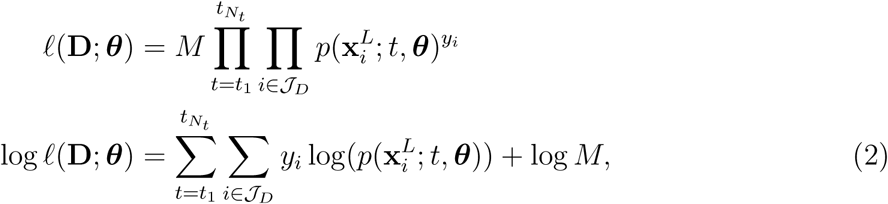

where *𝒥*_*D*_ is the set of states observed in the data, *M* is a combinatorial prefactor (i.e., from a multinomial distribution) that comes from the arbitrary reordering of measured data, and 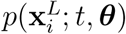 is the marginalized probability mass of the observable species,

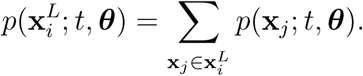

The vector of model parameters is denoted by ***θ*** = [*θ*_1_, *θ*_2_, …]. Neglecting the term log *M*, which is independent of the model, the summation in Eq. 2 can be rewritten as a product **y** log **p**^*L*^, where **y** ≡ [*y*_0_, *y*_1_, …] is the vector of the binned data, and 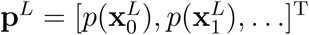 is the corresponding marginalized probability mass vector. One may then maximize Eq. 2 with respect to ***θ*** to find the *maximum likelihood estimate* (MLE) of the parameters, 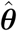, which will vary depending on each new set of experimental data. We next demonstrate how this likelihood function and the FSP model of the HOG1-MAPK induced gene expression system can be used to design optimal smFISH experiments using the FSP-based Fisher information matrix [6].

## RESULTS

### The Finite State Projection based Fisher information for models of signal-activated stochastic gene expression

The Fisher information matrix (FIM), is a common tool in engineering and statistics to estimate parameter uncertainties prior to collecting data, and which allows one to find experimental settings that can make these uncertainties as small as possible [3, 4, 31–34]. Recently, it has been applied to biological systems to estimate kinetic rate parameters in stochastic gene expression systems [3–6, 35]. In general, the FIM for a single measurement is defined:

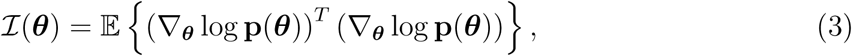

where the vector log **p**(***θ***) contains the log-probabilities of each potential observation, and the expectation is taken over the probability distribution of states **p**(***θ***) assuming the specific parameter set ***θ***. As the number of measurements, *N*_*c*_, is increased such that maximum likelihood estimates (MLE) of parameters are unbiased, the distribution of MLE estimates is known to approach a multivariate Gaussian distribution with a covariance given by the inverse of the FIM, i.e.,

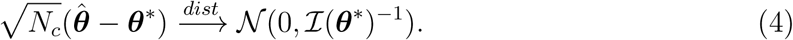

In [6], we developed the FSP-based Fisher information matrix (FSP-FIM), which allows one to use the FSP solution **p**(*t*), and its sensitivity 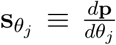, to find the FIM for stochastic gene expression systems. For a general FSP model, the dynamics of the sensitivity to each *j*^th^ kinetic parameter 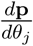 can be calculated according to:

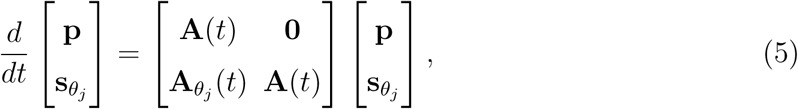

where 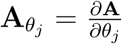. Solving Eq. 5 requires integrating a coupled set of ODEs that is twice as large as the original FSP system. The FSP-FIM at a single time *t* is then given by:

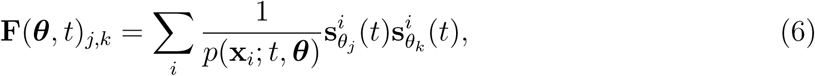

where the summation is taken over all states {**x**_*i*_} included in the FSP analysis (or over all observed states 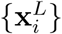 in the case of lumped observations). We note that the FSP computation of the FIM should be computationally tractable for problems for which the FSP solution itself is tractable. However, since the size of the FSP sensitivity matrix (Eq. 5) scales exponentially with the number of species, practical applications of the presented formulation of the FSP-FIM are currently restricted to models that have, or can be reduced to have, three or fewer distinct chemical species.

The FIM for a sequence of measurements taken independently (e.g., for smFISH data) at times 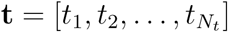 can be calculated as the sum across the measurement times:

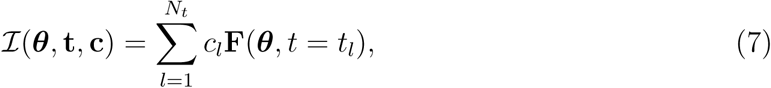

where 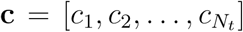 is the number of cells measured at each *l*^*th*^ measurement time. For smFISH experiments, the vector **c** plays an important role in the design of the study. By optimizing over all vectors **c** that sum to *N*_total_, one can find how many cells should be measured at each time point and which time points should be skipped entirely, (i.e., *c*_*l*_ = 0).

In the next subsection, we verify the FSP-FIM for this stochastic model with a time-varying parameter, and later find the optimal **c** for *STL1* mRNA in yeast cells.

### The FSP-FIM can quantify experimental information for stochastic gene expression under time-varying inputs

Our work in [6] was limited to models of stochastic gene expression that had piecewise constant reaction rates. Here, we extend this to time-varying reaction rates that affect the promoter switching in the system and which lead to time-varying **A**(*t*) in Eq. 5. For example, in the model depicted in Fig. 1a, the temporal addition of osmotic shock causes nuclear translocation of HOG1-MAPK, according to the time-varying function in Eq. 1.

Model parameters simultaneously fit to experimentally measured 0.2M and 0.4M *STL1* mRNA were adopted from [10] and used as a reference set of parameters (yellow dots in Fig. 2a and S1), which we define as ***θ***^*^. These reference parameters were used to generate 50 unique and independent simulated data sets, and each *n*^th^ simulated data set was fit to find the parameter set, 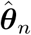, that maximizes the likelihood for that simulated data set. This process was repeated for two different experiment designs, including the original intuitive design from [10] (results shown in Fig. 2) and an optimized design discussed below (results shown in Fig. S1). To ease the computational burden of this fitting, the four parameters with the smallest sensitivities and largest uncertainties (i.e., those parameters that had the least effect on the model predictions and which were most difficult to identify) were fixed at their baseline values. The resulting MLE estimates for the remaining five parameters were collected into a set of 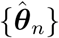 and are shown as yellow dots in Figs. 2 and S1. Using the asymptotic normality of the maximum likelihood estimator and its relationship to the FIM (Eq. 4), we then compared the 95% confidence intervals (CIs) of the inverse of the Fisher information (i.e., the Cramér Rao bound) to those of the MLE estimates (compare the purple and orange ellipses in Figs. 2a and S1a). We also compared the eigenvalues of the inverse of the Fisher information, {*v*_*i*_}, to the correspondingly ranked eigenvalues of the covariance matrix of MLE estimates, Σ_MLE_, in Figs. 2b and S1b. For further validation, we noted that the principle directions of the ellipses in Figs. 2a and S1a also match for the FIM and MLE analyses, as quantified by the angle between the paired FIM and Σ_MLE_ eigenvectors (Figs. 2b and S1b). For comparison, the angles between rank-matched eigenvectors of the FIM and Σ_MLE_ were all less than 12°, whereas non rank-matched eigenvectors were all greater than 79.9°. With the FSP-FIM verified for the HOG1-MAPK induced gene expression model, we next explore how the FSP-FIM can be used to optimally allocate the number of cells to measure at each time after osmotic shock.

**FIG. 2.**
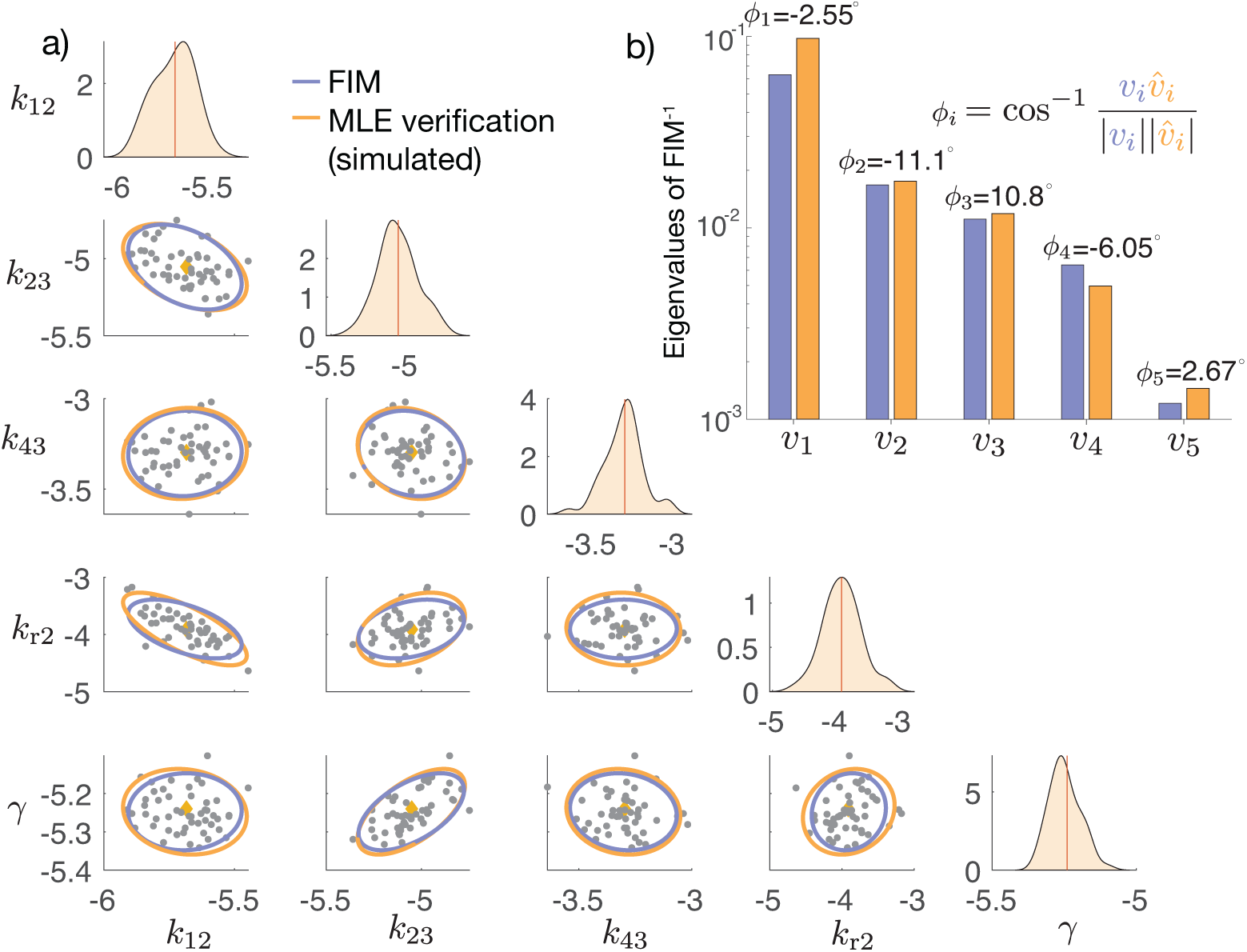
Verification of the FSP-FIM for the time-varying HOG1-MAPK model. (a) Marginal parameter histograms (top panels) and joint scatter plots (gray dots) for the MLE parameter estimates from 50 simulated data sets and for a subset of model parameters. All parameters are shown in logarithmic scale. The ellipses show the 95% CI for the inverse of the FIM (purple) and gaussian approximation of MLE scatter plot (orange). The yellow dots indicate the “true” parameters at which the FIM and simulated data sets were generated. (b) Rank-paired eigenvalues (*v*_*i*_) for the covariance of MLE estimates (orange) and inverse of the FIM (blue). The angles between corresponding rank-paired eigenvectors (*ϕ*_*i*_) are shown in degrees.

### Designing optimal measurements for the HOG1-MAPK pathway in *S. cerevisae*

To explore the use of the FSP-FIM for experiment design in a realistic context of MAPK-activated gene expression, we again utilize simulated time-course smFISH data for the osmotic stress response in yeast.

We start with a known set of underlying model parameters that were taken from simultaneous fits to 0.2M and 0.4M data in [10] (non-spatial model) to establish a baseline parameter set that is experimentally realistic. These parameters are then used to optimize the allocation of measurements at different time points *t* = [1, 2, 4, 6, 8, 10, 15, 20, 25, 30, 35, 40, 45, 50, 55] minutes after NaCl induction. Specifically, we ask what fraction of the total number of cells should be measured at each time to maximize the information about a specific subset of important model parameters. We use a specific experiment design objective criteria referred to as *D*_*s*_-optimality, which corresponds to minimizing the expected volume of the parameter space uncertainty for the specific parameters of interest [35], and which is found by maximizing the product of the eigenvalues of the FIM for those same parameters.

Mathematically, our goal is to find the optimal cell measurement allocation,

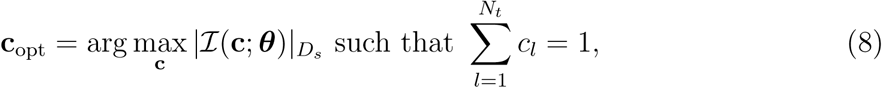

where *c*_*l*_ is the fraction of total measurements to be allocated at *t* = *t*_*l*_, and the metric 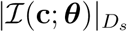 refers to the product of the eigenvalues for the total FIM (Eq. 7). The fraction of cells to be measured at each time point, **c**, was optimized using a greedy search, in which single-cell measurements were chosen one at a time according to which time point predicted the greatest improvement in the optimization criteria (see Supplementary Note 3 for more information).

To illustrate our approach, we first allocated cell measurements according to *D*_*s*_-optimality as found through this greedy search. Figure 3 shows the optimal fraction of cells to be measured at each time following a 0.2M NaCl input and compares these fractions to the experimentally measured number of cells from [10]. While each available time point was allocated a non-zero fraction of measurements, three time points at *t* = [10, 15, 30] minutes were vastly more informative than the other potential time points. To verify this result, we simulated 50 data sets of 1,000 cells each and found the MLE estimates for each sub-sampled data set. We compared the spread of these MLE estimates to the inverse of the optimized FIM, shown in Fig. S1.

**FIG. 3.**
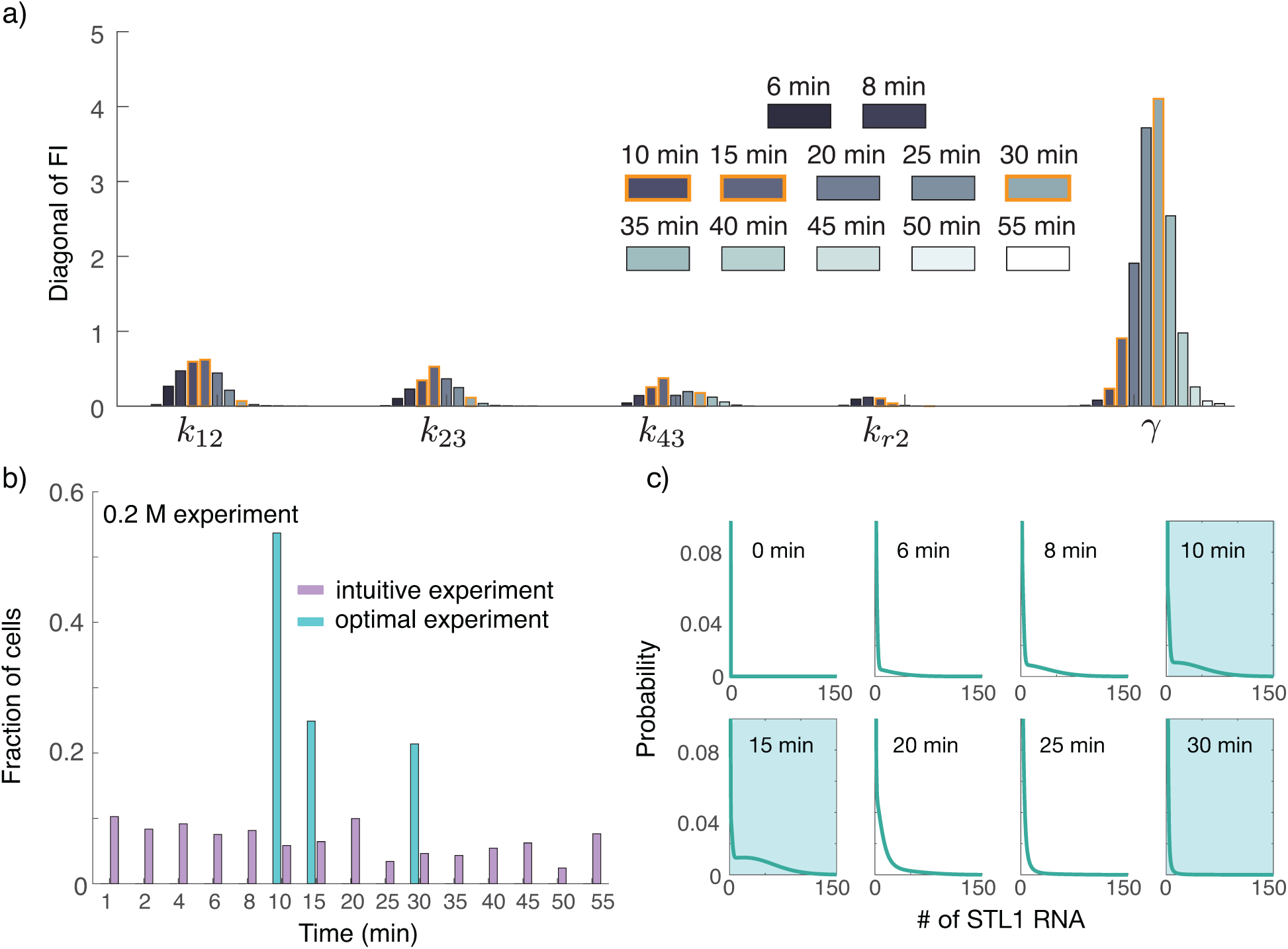
Optimizing the allocation of cell measurements at different time points. (a) Diagonal entries of the Fisher information at different measurement times. The optimal measurement times *t* = [10, 15, 30] minutes are highlighted in orange. (b) Comparison of optimal fractions of cells to measure (blue) at different time points determined by the FSP-FIM compared to experimentally measured numbers of cells at 0.2M NaCl (purple) from our work in [10]. (c) Probability distributions of *STL1* mRNA at several of measurement times. The blue boxes denote the time points of optimal measurements.

Comparing Figs. S1 with Fig. 2 illustrates the increase in information of the optimal 0.2M experiment compared to the intuitively designed experiment from [10]. In addition to providing much higher Fisher information, the optimal experiment requires measurement of only three time points compared to the 16 time points that were measured in the original experiment. Furthermore, we note that the FIM prediction of the MLE uncertainty is more accurate for the simpler optimal design, which is likely related to our observation that MLE estimates converge more easily for the optimized experiment design than they do for the original intuitive design.

Figure 4 next compares the *D*_*s*_-optimality criteria for the optimal (solid horizontal lines) and intuitive ([10], dashed horizontal lines) experiment designs to 1,000 randomly designed experiments for the 0.2M (black) and 0.4M (gray) conditions. To generate these random experiment designs, we selected a random subset of the measurement times, and allocated the total 1,000 cells among chosen time points using a multinomial distribution with equal probability for each time point. Figure 4a shows that the intuitive experiment is more informative than most random experiments, but is still substantially less informative than the optimal experiment.

**FIG. 4.**
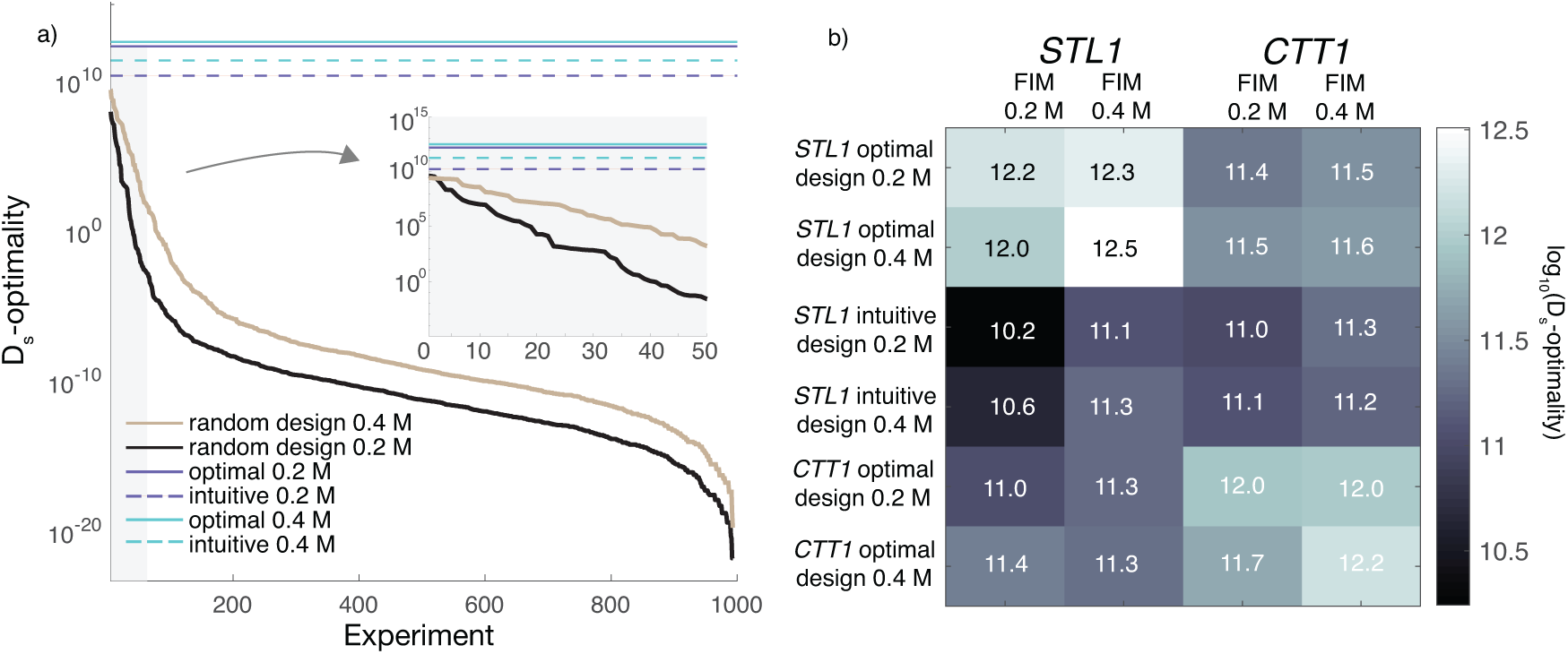
Information gained by performing optimal experiments compared to actual experiments. (a) *D*_*s*_-optimality for optimal design using three time points compared to the intuitive experiment designs made using 16 time points are shown with horizontal lines (purple, 0.2M and blue, 0.4M). Solid horizontal lines denote the optimal designs and dashed lines represent intuitive experiment designs. Randomly designed experiments with 0.2M and 0.4M NaCl are shown in black and orange. For the random experiments, the time points were selected by sampling them from the experimental measurement times, and then a random number of measurements were assigned to each selected time point. The inset shows the first 50 randomly designed experiments. (b) The *D*_*s*_-metric for different experiment designs (different rows) when applied to different genes or different experimental levels of osmotic shock (different columns). Lighter shades (higher *D*_*s*_-metrics) indicate experimental designs that are more suitable to identify parameters.

In many practical applications, a scientist would be unlikely to have precise *a priori* knowledge of model parameters prior to conducting experiments. Rather, they would have some estimate of these parameters, such as rough knowledge of appropriate time scales or existing data from another type of experiment. Such estimates could come from previous analyses of the system response to simpler experimental conditions, for measurements taken on slightly different cell lines or organisms, or considering results from different genes in related regulatory pathways. To explore the importance of knowing the exact process parameters or input dynamics prior to designing the experiment, we asked how well an experiment design optimized using parameters from one gene at a given level osmotic shock (e.g., *STL1* at 0.2M NaCl) would do to estimate parameters for another gene in a different osmotic shock condition (e.g., *CTT1* at 0.4M NaCl). Figure 4b demonstrates the impact of such mismatched experiment designs, where each row corresponds to a different intuitive or optimized experiment design (i.e., a specific allocation of cells to be measured at each time), and each column corresponds to a specific gene and specific osmotic shock condition to which that design could be applied. In all cases, the much simpler FIM-based optimal experiment designs perform as well or better than the more difficult intuitive designs, even when these FIM designs were computed assuming different environmental conditions and assuming genes whose parameters differ considerably from one another (see Supplemental Tables I and II for parameter sets). In other words, these results suggest that if one can compute a simple yet optimal experiment design based on one well-analyzed gene in a previously studied environmental condition, then that design may be equally valuable when applied to student a new, but related gene in a similar, yet slightly different context.

### Using the FSP-FIM to design optimal biosensor measurements

Thus far, and throughout our previous work in [6], we have sought to find the optimal set of experiments to reduce uncertainty in the estimates of *model parameters*. In this section, we discuss how the FSP-FIM allows for the optimization of experiment designs to address a more general problem of inferring *environmental variables* from cellular responses. Toward this end, we assume a known and parametrized model (i.e., the model defined above, which was identified previously in [10]), but which is now subject to unknown environmental influences. We explore what would be the optimal experimental measurements to take to characterize these influences. Specifically, we ask how many cells should be measured using smFISH, and at what times, to determine the specific concentration of NaCl to which the cells have been subjected – or, equivalently, we ask what experiments would be best suited to measure the effective stress induction level caused by addition of an unknown solution to the cells.

Recall from above that in the HOG1-MAPK transcription model, extracellular osmolarity ultimately affects stress response gene transcription levels through the time-varying parameter *k*_21_(*t*) (Eq. 1) as illustrated in Fig. 1b for 0.2M and 0.4M salt concentrations. Higher salt concentrations delay the time at which *k*_21_(*t*) returns to its nonzero value. The function in Eq. 1 can be coarsely approximated by the sum of three Heaviside step functions, *u*(*t* − *τ*_*i*_) as:

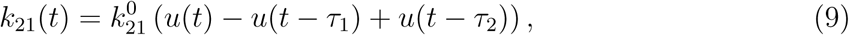

where *τ*_1_ is the fixed delay of the time it takes for nuclear kinase levels to reach the *k*_21_ deactivation threshold (about 1 minute or less, [9, 10]), and *τ*_2_ is the variable time it takes for the nuclear kinase to drop back below that threshold. In practice, the threshold-crossing time, *τ*_2_, should be directly related to the salt concentration experienced by the cell under reasonable salinity levels. This relationship is shown in Fig. 1b and 5b, where a 0.2M NaCl input exhibits a shorter *τ*_2_ than does a 0.4M input. For our analyses, we assume a prior uncertainty such that time *τ*_2_ can be any value uniformly distributed between 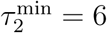 and 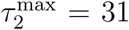 minutes, and our goal is to find the experiment that best reduces the posterior uncertainty in *τ*_2_ (and therefore could provide an estimate for the concentration of NaCl).

To reformulate the FSP-FIM to estimate uncertainty in *τ*_2_ given our model, the first step is to compute the sensitivity of the distribution of mRNA abundance to changes in the variable *τ*_2_ using Eq. 5, in which 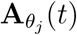 is replaced with 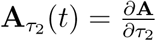 as follows:

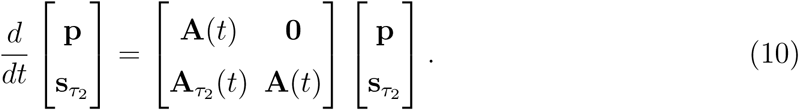

As *k*_21_(*t*) is the only parameter in **A** that depends explicitly on *τ*_2_, all entries of 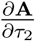 are zero except for those which depend on *k*_21_(*t*), and

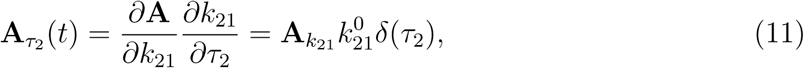

and therefore 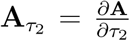 is non-zero only at *t* = *τ*_2_. Using this fact, the equation for the sensitivity dynamics is uncoupled from the FSP dynamics for *t* ≠ *τ*_2_, and can be written simply as:

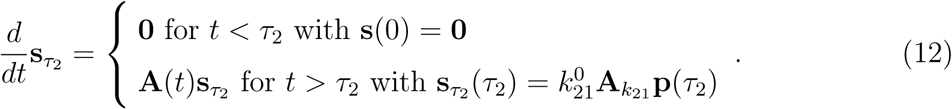

If the Fisher information at each measurement time is written into a vector 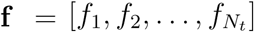 (noting that the Fisher information at any time *t*_*l*_ is the scalar quantity, *f*_*l*_), and the number of measurements per time point is the vector, 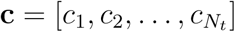, then the total information for a given value of *τ*_2_ can be computed as the dot product of these two vectors,

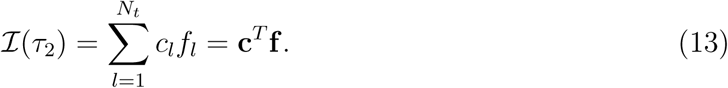

Our goal is to find an experiment that is optimal to determine the value of *τ*_2_, given an assumed prior that *τ*_2_ is sampled from a uniform distribution between 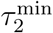 and 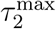. To find the experiment **c**_opt_ that will reduce our posterior uncertainty in *τ*_2_, we integrate the inverse of the FIM in Eq. 13 over the prior uncertainty in *τ*_2_,

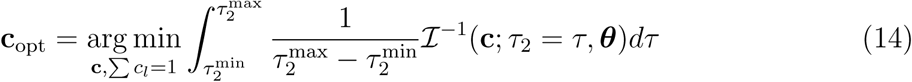

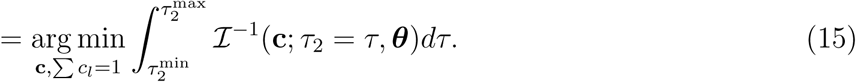

For later convenience, we define the integral in Eq. 14 (i.e., the objective function of the minimization) by the symbol *𝒥*, which corresponds to the expected uncertainty about the value of *τ*_2_ for a given **c**.

Next, we apply the greedy search from above to solve the minimization problem in Eq. 15 to find the experiment design **c**_opt_ that minimizes the estimation error of *τ*_2_. Figure 6 shows examples of seven different experiments to accomplish this task, ranked according to the FSP-FIM value *𝒥* from most informative (top left) to least informative (bottom left), but all using the same number of measured cells. For each experiment, the FSP-FIM was used to estimate the posterior uncertainty (i.e., expected standard deviation) in the estimation of *τ*_2_, which is shown by the orange bars in Fig. 6. To verify these estimates, we then chose 64 uniformly spaced values of *τ*_2_, which we denote as the set 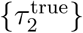, and for each 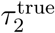, we simulated 50 random data sets of 1,000 cells distributed according to the specified experiment designs. For each of the 64×50 simulated data sets, we then determined the value 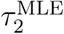 between 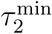 and 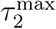 that maximized the likelihood of the simulated data according to Eq. 2. The root mean squared estimate (RMSE) error over all random values of 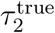 and estimates, 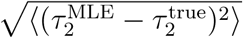, was then computed for each of the six different experiment designs. Figure 6 shows that the FIM-based estimation of uncertainty and the actual MLE-based uncertainty are in excellent agreement for all experiments (compare purple and orange bars). Moreover, it is clear that the optimal design selected by the FIM-analysis performed much better to estimate *τ*_2_ than did the uniform or random experimental designs. A slightly simplified design, which uses the same time points as the optimal, but with equal numbers of measurements at each time, performed nearly as well as the optimal design.

**FIG. 5.**
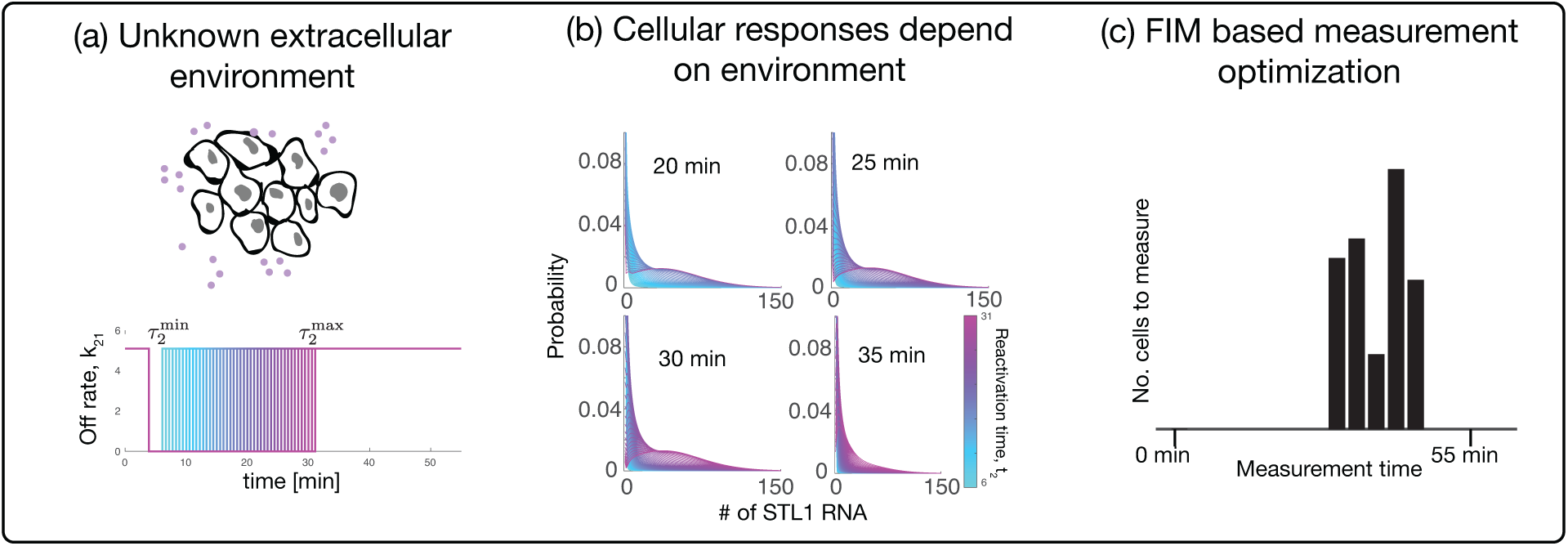
Overview of optimal design for biosensing experiments for the osmotic stress response in yeast. (a) Unknown salt concentrations (purple dots) in the environment give rise to different reactivation times, *τ*_2_, which affect the gene expression in the model through the rate *k*_21_. These different reactivation times cause downstream *STL1* expression dynamics to behave differently as shown in panel (b). (c) Different responses can be used to resolve experiments that reduce the uncertainty in *τ*_2_.

**FIG. 6.**
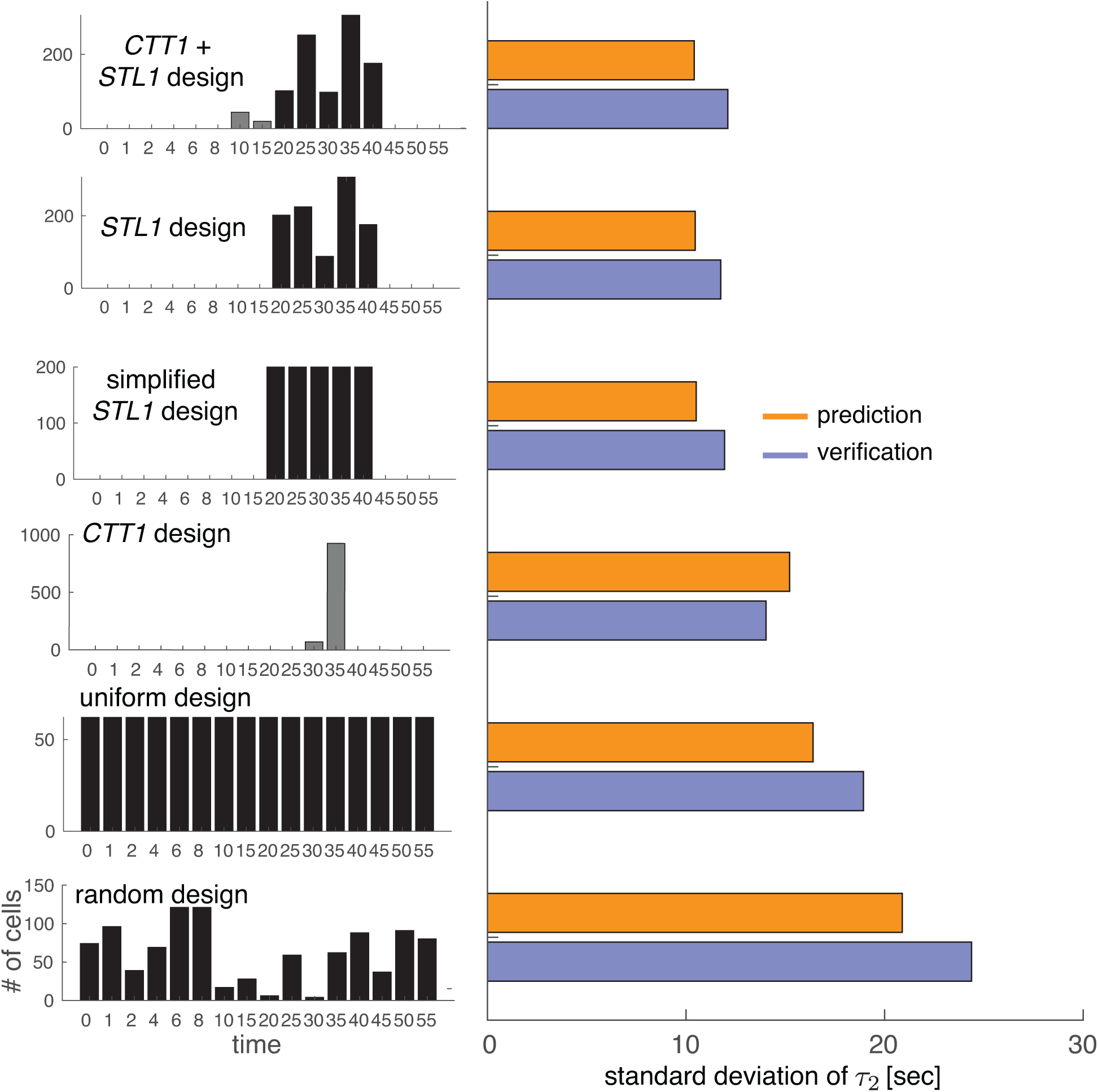
Verification of the uncertainty in τ_2_ for different experiment designs. The left panel shows various experiment designs, where the sum of the bars (i.e., the total number of measurements) is 1,000. Gray bars represent the measurements of *CTT1* and black bars *STL1*. The right panel shows the value of the objective function in Eq. 14 for each experiment design in orange, and the RMSE values for verification are shown in purple.

The set of experiment designs shown in Fig. 6 includes the best design that only uses *STL1* (second from top), the best design that uses only *CTT1* (fourth from top), and the best designs that uses some cells with *CTT1* and some with *STL1* (top design). To find the best experiment design for measurement of two different genes, we assumed that at each time, either *STL1* mRNA *or CTT1* mRNA (but not both) could be measured, corresponding to using smFISH oligonucleotides for either *STL1* or *CTT1*. To determine which gene should be measured at each time, we compute the Fisher information for *CTT1* and *STL1* for every measurement time and averaged this value over the range of *τ*_2_. For each measurement time *t*_*l*_, the gene is selected that has the higher average Fisher information for *τ*_2_. The number of cells per measurement time were then optimized as before, except the choice to measure *CTT1* or *STL1* was based on which mRNA had the larger Fisher information (Eq. 13) at that specific point in time. The best *STL1*-only experiment design was found to yield uncertainty of 10.5 seconds (standard deviation); the best *CTT1*-only experiment was found to yield an uncertainty of 15.2 seconds and the best mixed *STL1*/*CTT1* experiment design was found to yield an uncertainty of 10.4 seconds. In other words, for this case the *STL1* gene was found to be much more informative of the environmental condition than was *CTT1*, and the use of both *STL1* and *CTT1* provides only minimal improvement beyond the use of *STL1* alone. We note that although measurement times in the optimized experiment design were restricted to a resolution of five minutes or more, the value of *τ*_2_ could be estimated with an error of only 10 seconds, corresponding to a roughly 30-fold improvement of temporal resolution beyond the allowable sampling rate.

### Experimental validation for FSP-FIM based designs of biosensor measurements

To experimentally validate our FSP-FIM based approach to design optimal measurement times, we next examined experimental smFISH data taken for the *STL1* and *CTT1* genes at different times following yeast osmotic shock [10]. These data include a total of 535-4808 cells measured at each of 16 time points following osmotic shocks of 0.2M or 0.4M NaCl. We asked how well could we identify the concentration of the osmotic shock from the experimental data using only 75 individual cells per experiment. We again proposed the six different potential experiments depicted in Fig. 6, including: the optimal *STL1* and *CTT1* design, the optimal *STL1* design, the simplified *STL1* design with 15 cells for each of the optimal five time points, the optimal *CTT1* design, the uniform *STL1* design, and the random STL1 design. For each design, we created 1,000 different experimental replica datasets, each consisting of 100 cells randomly chosen from the original data. For each replica data set, we then used the CME model (Supplementary Note 1) with a parametrized form of the HOG1-MAPK nuclear localization signal (Supplementary Note 2) to find the NaCl concentration that maximizes the likelihood of the data given the model.

Figure 7 shows the resulting histograms for the estimated NaCl concentrations for each of the six experiment designs, when the cells were actually subjected to experimental osmotic shocks of 0.2M NaCl (Fig. 7a) or 0.4M NaCl (Fig. 7c). From Figures 7a,c, it is clear that the FSP analysis provides an accurate estimate for the level of the osmotic shock input using a relatively small number of cells, despite the fact that producing such estimates was not an intended use of the model in its original formulation or parameter inference [9, 10]. Figures 7b,d show the uncertainty (standard deviation) in the experimental estimate of NaCl concentration (light bars), when cells are collected according to the six specific experiment designs, and compares these results to the FSP-FIM uncertainty estimates (dark bars) using the simplified step input function (Eq. 9). With the exception of the sub-optimal *CTT1*-only design, the close matches between the relative trends of the variance in experimental estimation of NaCl and the variance predicted by the FSP-FIM analysis with the approximated step-function input gives further experimental validation that the FSP-FIM approach can be used to choose more informative experiment designs, even in cases where the FSP analyses uses inexact assumptions for model kinetics. The single discrepancy in trends led us to more closely examine the model and experimental data for *CTT1* expression at the 35 minute time point that dominates the *CTT1*-only design. By examining Supplemental Figure S7 from [10], we found that this specific combination of *CTT1* at 35 minutes following 0.4M NaCl osmotic shock showed a greater discrepancy between model and data than any of the other 63 combinations of 16 times, two genes and two conditions, yet it is unclear if that difference was an artifact of the experiment or an actual transient effect that only affected that specific combination of gene, time, and environmental condition.

**FIG. 7.**
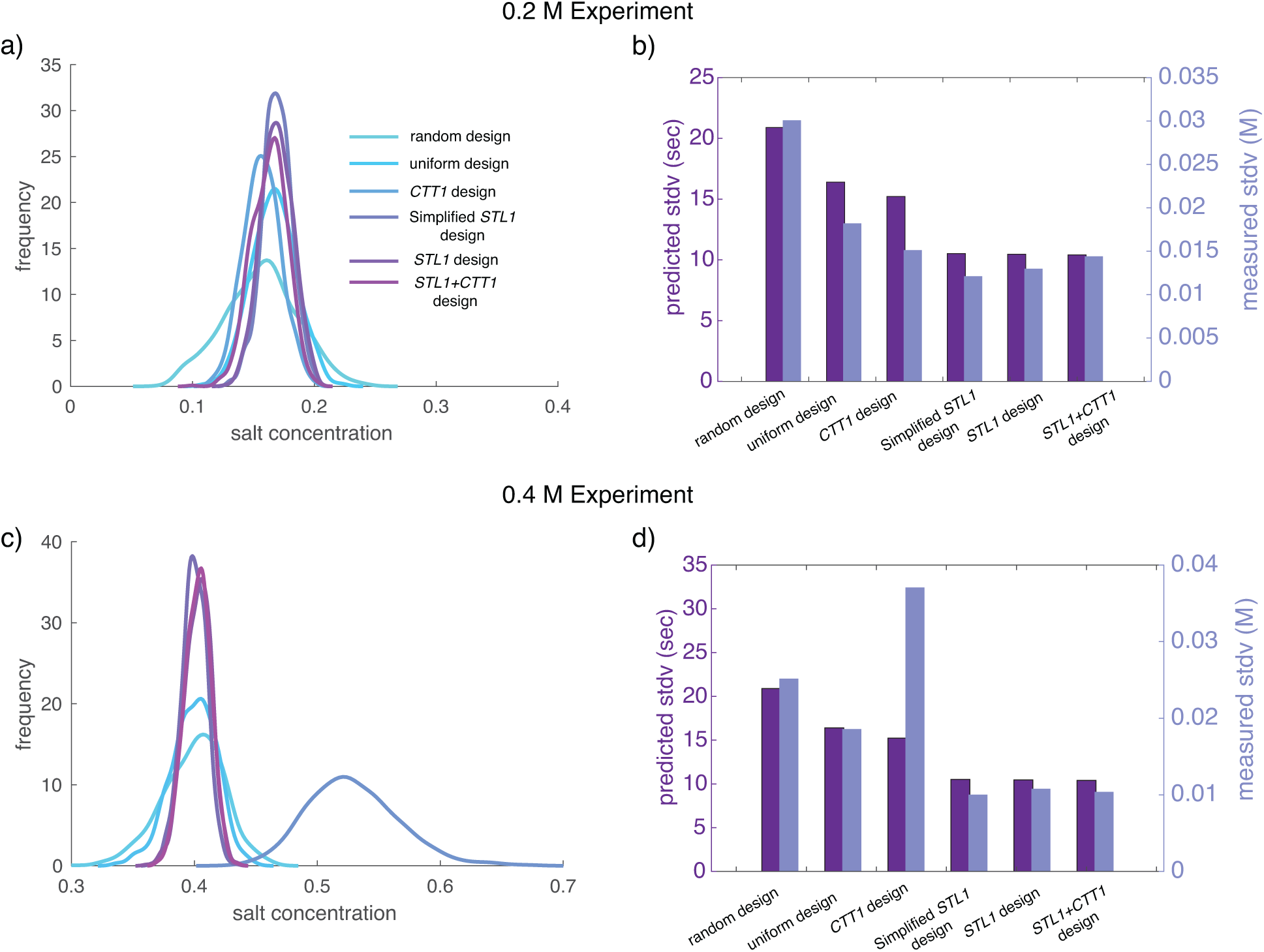
Experimental validation of FSP-FIM based design for optimal biosensor measurements. (a) Distribution of FSP-based MLE estimates for NaCl concentration using the six experimental designs from Fig 6. Each distribution comes from 1,000 replicas of 75 cells per replica spread out over the possible 16 time points. Replica data were sampled randomly from published experimental data [10] that contain two or three biological replicas and 535-4808 cells per time point. The true experimentally applied level of osmotic shock was 0.2M NaCl. (b) The MLE estimation standard deviation for each experiment design applied to a data set taken at 0.2M NaCl (blue). These deviations are compared to FSP-FIM deviation predictions using a piecewise constant model for HOG1 nuclear localization (purple). (c,d) Same as (a,b) but for a true NaCl concentration of 0.4M.

## DISCUSSION

The methods developed in this work present a principled, model-driven approach to allocate how many snapshot single-cell measurements should be taken at each time during analysis of a time-varying stochastic gene regulation system. We demonstrate and verify these theories on a well-established model of osmotic stress response in yeast cells, which is activated upon the nuclear localization of phosphorylated HOG1 [9, 10]. For this system, we showed how to optimally allocate the number of cells measured at each time so as to maximize the information about a subset of model parameters. We found that the optimal experiment design to estimate model parameters for the *STL1* gene only required three time points. Moreover, these three time points (*t* = [10, 15, 30] minutes, highlighted by blue in Fig. 3b) are at biologically meaningful time points. At *t* = 10 and 15 minutes, the system is increasing to maximal expression, and the probability to measure a cell with elevated mRNA content is high, which helps reduce uncertainty about the parameters in the model that control maximal expression. Similarly, at the final experiment time of *t* = 30 minutes, the system is starting to shut down gene expression, and therefore this time is valuable to learn about the time scale of deactivation in the system as well as the mRNA degradation rate. These effects are clearly illustrated in Fig. 3a, which shows that times *t* = 10 and *t* = 15 minutes provide the most information about parameters *k*_12_, *k*_23_ and *k*_43_, whereas measurements at *t* = 30 minutes provide the most information about *γ*. Because *γ* is the easiest parameter to estimate (e.g., its information is greater), not as many cells are needed at *t* = 30 minutes to constrain that parameter. Similarly, because *k*_*r*2_ is the most difficult parameter to estimate (e.g., it has the lowest information across all experiments), and because *t* = 10 minutes is one of the few time points to provide information about *k*_*r*2_, the optimal experimental design selects a large number of cells at the time *t* = 10 minutes. This analysis demonstrates that the optimal experiment design can change depending upon which parameters are most important to determine (e.g., *γ* or *k*_*r*2_ in this case), a fact that we expect will be important to consider in future experiment designs.

Because we constrained all potential experiment designs to be within the subset of experiments performed in our previous work [10], we are able to compare the information of optimal experiment designs to intuitive designs that have actually been performed. We found that while the intuitive experiments were almost always better than could be expected by random chance, they still provided several orders of magnitude lower Fisher information than would be possible with optimal experiments (Fig. 4a). Moreover, in our analyses, we found that optimal designs could require far fewer time points than those designed by intuition (e.g., only three time points were needed in Fig. 3), and therefore these designs can be much easier and less expensive to conduct. We also found that utility of optimal experiment designs could be relatively insensitive to variation in the experimental conditions or the specific model parameters used for the experiment design. For example, we found that experiments optimized for one gene at one level of osmotic shock were still at least as good–and in most cases better–than intuitive designs, even when conducted using different genes and at a different level of osmotic shock (Fig. 4b). In practice, this fact would allow for effective experiment designs despite inaccurate prior assumptions.

In addition to suggesting optimal experiments to identify model parameters, we showed that the FSP approach could be used to infer parameters of fluctuating extracellular environments from single-cell data and that the FSP-FIM combined with an existing model could be used to design optimal experiments to improve this inference (Figs. 5 and 6). We experimentally verified this potential by examining many small sets of single-cell smFISH measurements for different genes and different measurement times, and we showed that an FSP-FIM analysis could correctly rank which experiment designs would give the best estimates of osmotic shock environmental conditions. Along a very similar line of reasoning, one can also adapt the FSP-FIM analysis to learn what biological design parameters would be optimal to reduce uncertainty in the estimate of important environmental variables. For example, Fig. 8 shows the expected uncertainty in *τ*_2_ as a function of the degradation rate of the *STL1* gene assuming that 50 cells could be measured at each experimental measurement time *t* = [1, 2, 4, 6, 8, 10, 15, 20, 25, 30, 35, 40, 45, 50, 55] minutes using the smFISH approach. We found that the best choice for *STL1* degradation rate to most accurately determine the extracellular fluctuations would be 2.4 × 10^−3^ mRNA/min, which is about half of the experimentally determined value of 5.3 × 10^−3^ ± 5.9 × 10^−5^ from [10]. This result is consistent with our earlier finding that the faster degrading *STL1* mRNA is a much better determinant of the HOG1 dynamics than is the slower-degrading *CTT1* mRNA, and suggests that other less stable mRNA could be more effective still. We expect that similar, future applications of the FSP-based Fisher information will be valuable in other systems and synthetic biology contexts where scientists seek to explore how different cellular properties affect the transmission of information between cells or from cells to human observers. Indeed, similar ideas have been explored recently using classical information theory in [36–39], and recent work in [7, 40] has noted the close relationship between Fisher information and the channel capacity of biochemical signaling networks.

**FIG. 8.**
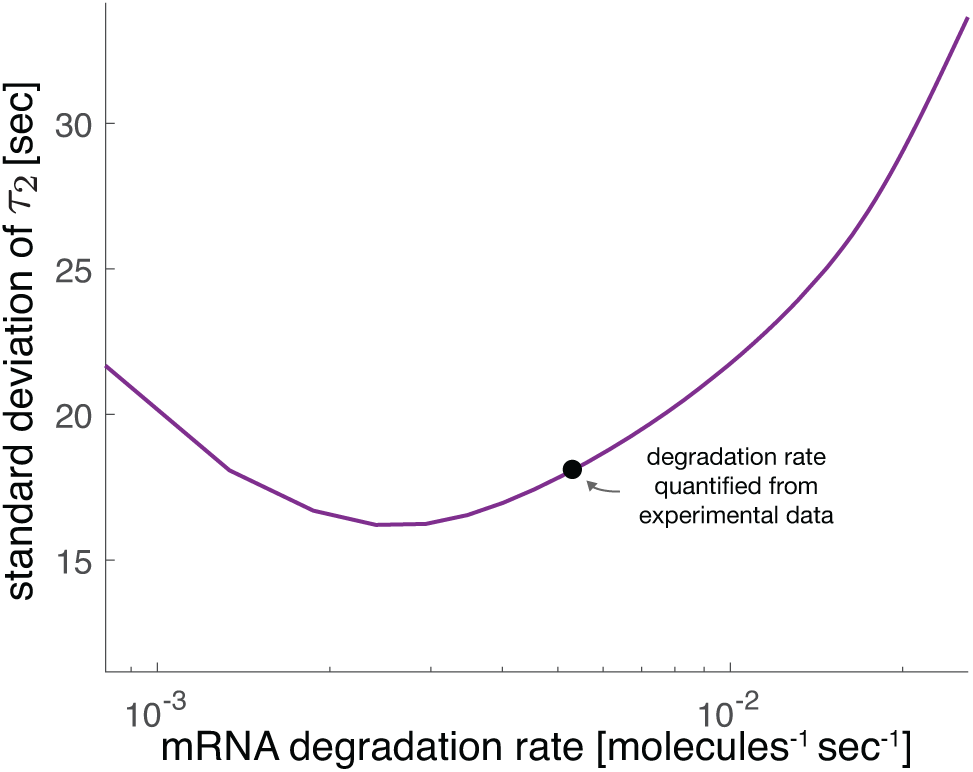
Optimal mRNA degradation rates to reduce uncertainty about the extracellular environment. Uncertainty in the time at which the *STL1* gene turns off, *τ*_2_, as a function of mRNA degradation rate (purple). The black dot corresponds to the degradation rate that was quantified from experimental data.

We expect that computing optimal experiment designs for time-varying stochastic gene expression will create opportunities that could extend well beyond the examples presented in this work. Modern experimental systems are making it much easier for scientists and engineers to precisely perturb cellular environments using chemical induction [41–43] or optogenetic control [44–46]. Many such experiments involve stochastic bursting behaviors at the mRNA or protein level [8–10, 45], and precise optimal experiment design will be crucial to understand the properties of stochastic variations in such systems. A related field that is also likely to benefit from such approaches is biomolecular image processing and feedback control, for which one may need to decide in real time which measurements to make and in what conditions.

## CONTENTS OF SUPPLEMENTAL INFORMATION

- **Supplementary Note 1**: Detailed description of the FSP model for stress response genes in yeast. This section includes an analysis of sensitivities of the model to different kinetic parameters.
- **Supplementary Note 2**: Discussion of the the HOG1-MAPK nuclear localization model and how it was approximated for the section of the manuscript on optimal biosensors.
- **Supplementary Note 3**: Description of the algorithm used to find optimal experiments.
- **Supplementary Table 1**: Parameters for the stochastic model for the STL1 and CTT1 genes.
- **Supplementary Table 2**: HOG-Signaling Model Parameters for 0.4M and 0.2M experimental conditions.
- **Supplementary Figure 1**: Verification of the FSP-FIM for the time-varying HOG1-MAPK model with the optimal experiment design.

## DATA AVAILABILITY

All data and codes associated with this article will be made available upon acceptance of the article at: https://github.com/MunskyGroup/fox_et_al_complexity_2019.

## ACKNOWLEDGEMENTS

ZRF and BEM were supported by National Institutes of Health [R35 GM124747]. ZRF was also supported by the Agence Nationale de la Recherche [ANR-18-CE91-0002, CyberCircuits]. GN was supported by National Institutes of Health [DP2 GM11484901, R01GM115892] and Vanderbilt Startup Funds. The presented analyses used the computational resources of the W M Keck High Performance Compute Cluster supported under a W M Keck Foundation Award. The content is solely the responsibility of the authors and does not necessarily represent the official views of the funding agencies.

