## Supplementary Materials for "Optimal Design of Single-Cell Experiments within Temporally Fluctuating Environments"

### Supplementary information for: Optimal Allocation of Single-Cell Measurements for the HOG-MAPK Pathway in S. Cerevisae

Zachary R Fox

Gregor Neuert

*Department of Molecular Physiology and Biophysics,*

*School of Medicine, Vanderbilt University, Nashville, TN 37232, USA*

*Department of Biomedical Engineering, School of Engineering,*

*Vanderbilt University, Nashville, TN 37232, USA and*

*Department of Pharmacology, School of Medicine,*

*Vanderbilt University, Nashville, TN 37232, USA*

Brian Munsky

*Department of Chemical and Biological Engineering,*

*Colorado State University Fort Collins, CO 80523, USA and*

*School of Biomedical Engineering, Colorado State University Fort Collins, CO 80523, USA*

#### SUPPLEMENTARY NOTE 1: STOCHASTIC MODEL OF YEAST STRESS RESPONSE

The chemical master equation is the workhorse of stochastic systems biology. It describes the time evolution of the probabilities of the system states $p(\mathbf{x}, t)$, where $\mathbf{x}_i = [\eta_1, \eta_2, \ldots, \eta_M]$ describes the integer counts of the number of molecules in the system [1]. The probability, sometimes called propensity, to transition between these states in the infinitesimal time $dt$ is given by $w(x, t, \boldsymbol{\theta})dt$. Biochemical systems are often described in terms of chemical reactions, in which molecules like DNA, RNA, and proteins are created, destroyed, diluted, or modified. From the theory of biochemical kinetics, these reaction rates can be taken as propensities of a given reaction in the system occuring. Each propensity function is associated with a corresponding stoichiometry vector $\boldsymbol{\psi}_\nu$, which describes how the state $\mathbf{x}$ changes when a given the $\nu^{\text{th}}$ reaction occurs. Thus, the chemical master equation is given by

$$\frac{dp(\mathbf{x}_i, t; \boldsymbol{\theta})}{dt} = -\sum_{\nu=1}^{R} p(\mathbf{x}_i, t; \boldsymbol{\theta}) w_\nu(\mathbf{x}_i, t; \boldsymbol{\theta}) + \sum_{\nu=1}^{R} p(\mathbf{x}_i - \boldsymbol{\psi}_\nu, t; \boldsymbol{\theta}) w_\nu(\mathbf{x}_i - \boldsymbol{\psi}_\nu, t; \boldsymbol{\theta}). \quad (1)$$

Often, the probability vector is combined over all the states $\mathbf{p} = [p(\mathbf{x}_i)]$, and therefore Eq. 1 is written in linear form as $\frac{d\mathbf{p}}{dt} = \mathbf{A}\mathbf{p}$. However, this set of ODEs often has infinite dimension, and therefore cannot be easily solved. The finite state projection approach to solving the chemical master equation approximates the infinite state Markov chain in Eq. 1 by truncating it into a finite subset of linear ODEs [2]. This is achieved by first partitioning the entire state space of the system $\mathbf{X}$ into two disjoint sets: a finite set of states $\mathbf{X}_{\mathcal{J}}$ and an infinite set of states $\mathbf{X}_{\mathcal{J}'}$. The FSP approach then combines all of $\mathbf{X}_{\mathcal{J}'}$ into a single state $g(t)$, and does not allow any reactions to leave this space. The dynamics for the FSP system are then given by the finite, linear set of ODEs,

$$\frac{d}{dt}\begin{bmatrix} \mathbf{p}_{\mathcal{J}}(t; \boldsymbol{\theta}) \\ g(t; \boldsymbol{\theta}) \end{bmatrix} \begin{bmatrix} \mathbf{A}_{\mathcal{J}\mathcal{J}} & 0 \\ -\mathbf{1}^{\mathrm{T}}\mathbf{A}_{\mathcal{J}\mathcal{J}} & 0 \end{bmatrix} \begin{bmatrix} \mathbf{p}_{\mathcal{J}}(t; \boldsymbol{\theta}) \\ g(t; \boldsymbol{\theta}) \end{bmatrix}, \quad (2)$$

where $\mathbf{A}_{\mathcal{J}\mathcal{J}}$ is the finite submatrix taken from $\mathbf{A}$ according the the states of the CME which were retained $\mathbf{X}_J$. For more details on the FSP approach, we refer the reader to [2–5]. For STL1 and CTT1 gene expression in yeast, the FSP matrices can be written as a block diagonal matrix,

$$\frac{d}{dt}\begin{bmatrix}\mathbf{p}_0\\ \mathbf{p}_1\\ \mathbf{p}_2\\ \vdots\\ \mathbf{P}_{N_m}\\ g(t)\end{bmatrix}=\begin{bmatrix}\mathbf{S}-\mathbf{T} & \mathbf{\Gamma} & \mathbf{0} & \dots & 0\\ \mathbf{T} & \mathbf{S}-\mathbf{T}-\mathbf{\Gamma} & 2\mathbf{\Gamma} & \ddots & 0\\ \mathbf{0} & \mathbf{T} & \mathbf{S}-\mathbf{T}-2\mathbf{\Gamma} & \ddots & 0\\ \vdots & \ddots & \ddots & \mathrm{N_m}\mathbf{\Gamma} & 0\\ \mathbf{0} & \dots & \mathbf{T} & \mathbf{S}-\mathbf{T}-\mathrm{N_m}\mathbf{\Gamma} & 0\\ \mathbf{0} & \dots & \dots & \mathbf{1}^T\mathbf{T} & 0\end{bmatrix}\begin{bmatrix}\mathbf{p}_0\\ \mathbf{p}_1\\ \mathbf{p}_2\\ \vdots\\ \mathbf{P}_{N_m}\\ g(t)\end{bmatrix}, \tag{3}$$

where the block matrices $\mathbf{S}$, $\mathbf{T}$, and $\mathbf{\Gamma}$ are given by:

$$\mathbf{S}(t)=\begin{bmatrix}-k_{12} & k_{21}(t) & 0 & 0\\ k_{12} & -k_{21}(t)-k_{23} & k_{32} & 0\\ 0 & k_{23} & -k_{32}-k_{34} & k_{43}\\ 0 & 0 & k_{34} & -k_{43}\end{bmatrix};$$
$$\mathbf{T}=\begin{bmatrix}k_{r_1} & 0 & 0 & 0\\ 0 & k_{r_2} & 0 & 0\\ 0 & 0 & k_{r_3} & 0\\ 0 & 0 & 0 & k_{r_4}\end{bmatrix};$$
$$\mathbf{\Gamma}=\begin{bmatrix}\gamma & 0 & 0 & 0\\ 0 & \gamma & 0 & 0\\ 0 & 0 & \gamma & 0\\ 0 & 0 & 0 & \gamma\end{bmatrix}. \tag{4}$$

.

Note that each submatrix $\mathbf{S}$, $\mathbf{T}$, $\mathbf{\Gamma}$ can be futher decomposed into sub-sub matrices of 1's and 0's multiplied by a single model parameter. For example,

$$\mathbf{T}=k_{\mathrm{r1}}\begin{bmatrix}1&0&0&0\\0&0&0&0\\0&0&0&0\\0&0&0&0\end{bmatrix}+k_{\mathrm{r2}}\begin{bmatrix}0&0&0&0\\0&1&0&0\\0&0&0&0\\0&0&0&0\end{bmatrix}+k_{\mathrm{r3}}\begin{bmatrix}0&0&0&0\\0&0&0&0\\0&0&1&0\\0&0&0&0\end{bmatrix}+k_{\mathrm{r4}}\begin{bmatrix}0&0&0&0\\0&0&0&0\\0&0&0&0\\0&0&0&1\end{bmatrix}. \tag{5}$$

All master equations which are linear in parameters can be decomposed like this, and therefore the generator the entire generator matrix can easily be constructed as the sum of each matrix,

$$\mathbf{A} = \sum_{i=1}^{N_{\boldsymbol{\theta}}} \boldsymbol{\theta}_i A_{\boldsymbol{\theta}_i}. \tag{6}$$

Furthermore, the sensitivity matrices $\frac{\partial \mathbf{A}}{\partial \boldsymbol{\theta}_i} \equiv \mathbf{A}_{\boldsymbol{\theta}_i}$ required to compute the FSP-FIM can easily be found from the decomposed matrices in Eq. 5, i.e.

$$\mathbf{T}_{k_{r1}} = \begin{bmatrix} 1 & 0 & 0 & 0 \\ 0 & 0 & 0 & 0 \\ 0 & 0 & 0 & 0 \\ 0 & 0 & 0 & 0 \end{bmatrix} \tag{7}$$

and

$$\mathbf{A}_{k_{r1}} = \begin{bmatrix} -\mathbf{T}_{k_{r1}} & \mathbf{0} & \mathbf{0} & \dots & \mathbf{0} \\ \mathbf{T}_{k_{r1}} & -\mathbf{T}_{k_{r1}} & \mathbf{0} & \ddots & \mathbf{0} \\ \mathbf{0} & \mathbf{T}_{k_{r1}} & -\mathbf{T}_{k_{r1}} & \ddots & \mathbf{0} \\ \vdots & \ddots & \ddots & \ddots & \vdots \\ \mathbf{0} & \dots & \dots & \mathbf{T}_{k_{r1}} & -\mathbf{T}_{k_{r1}} \end{bmatrix} \tag{8}$$

In [5], we derive the sensitivity dynamics to parameter $\boldsymbol{\theta}_i$ as

$$\frac{d}{dt} \begin{bmatrix} \mathbf{p} \\ \mathbf{s}_{\boldsymbol{\theta}_i} \end{bmatrix} = \begin{bmatrix} \mathbf{A} & \mathbf{0} \\ \mathbf{A}_{\boldsymbol{\theta}_i} & \mathbf{A} \end{bmatrix} \begin{bmatrix} \mathbf{p} \\ \mathbf{s}_{\boldsymbol{\theta}_i} \end{bmatrix}, \tag{9}$$

which are then used to compute the Fisher information matrices, as described in the main text.

#### SUPPLEMENTARY NOTE 2: NUCLEAR LOCALIZATION OF HOG-MAPK

The nuclear localization of HOG-MAPK has been studied in significant detail [6–9]. In [10, 11], we modeled the nuclear enrichment of HOG-MAPK, $f(t)$ with the empirical equation,

$$f(t) = A \left( \frac{(1 - \mathrm{e}^{-r_1 t}) e^{-r_2(S)t}}{1 + \frac{(1-\mathrm{e}^{-r_1 t}) e^{-r_2(S)t}}{m}} \right)^{\eta}, \tag{10}$$

as shown in Fig. 1b in the main text, where $S$ is the NaCl concentration and

$$r_2(S) = \frac{\alpha}{S - \Delta}. \tag{11}$$

Model parameters are given in Table I. In [10], these parameters where fit to experimentally measured localization levels at 0.2 M and 0.4 M NaCl concentrations. As noted in Eq. 1 of the main manuscript, these nuclear MAPK levels are then mapped to a thresholded reactivation function,

$$k_{21}(t) = \max\left[0, \alpha - \beta f(t)\right]. \tag{12}$$

To illustrate how the osmotic shock reponse model can be used to reduce uncertainty in the model, we connect the environmental salinity level to the reactivation time $\tau_2$, shown in Fig. 5 in the main manuscript. To find the reactivation times, we assumed that the salinities are linearly related to the reactivation times, that is $\tau_2 = cM$, where $c$ is the coefficient of proportionality. We used reactivation times determined from the model in [11] for 0.2M and 0.4M experimental conditions to find $c$, and in turn could estimate reactivation times for reasonable salinity levels. We then use these estimates of reactivation times in our reduced piecewise constant model, Eq. 9 in the main text. We note that this choice of model for reactivation times (i.e. a line) could be made more precise with more experimental data or a secondary model for HOG-MAPK nuclear localization.

#### SUPPLEMENTARY NOTE 3: OPTIMIZATION OF CELL MEASUREMENTS

To calculate the optimal sampling schedule for the FIM, we want to find the vector $\mathbf{c}$ which maximizes the FIM (see Eq. 8 in the main text), in terms of $D_s$ optimality, which corresponds to maximizing the product of the eigenvalues of the FIM. To optimize the number of measurements to be taken at a given time, we use a ‘greedy’ optimization approach, described in Algorithm 1.

**Algorithm 1** Measurement allocation optimization algorithm.

Initialize $\mathbf{c}$

Initialize $\mathbf{f}$

$i = 0$

**while** $\mathbf{c}^i(\mathbf{c}^i\mathbf{1})^{-1} \not\approx \mathbf{c}^{i-1}(\mathbf{c}^{i-1}\mathbf{1})^{-1}$ **do**

  $\hat{\mathbf{c}} = \mathbf{c}$

  **for** $k \in (1, N_t)$ **do**

    $\hat{c}_k = c_k + 1$

    $q_k = \hat{\mathbf{c}}^{\mathbf{T}}\mathbf{f}$

  **end for**

  $j = \text{argmax}(\mathbf{q})$

  $c_j^i = c_j^i + 1$

  $i = i + 1$

**end while**

#### SUPPLEMENTARY TABLES

TABLE I. HOG-MAPK Model Parameters

| gene | $k_{12}$ $[s^{-1}]$ | $\alpha$ $[s^{-1}]$ | $\beta$ | $k_{23}$ $[s^{-1}]$ | $k_{32}$ $[s^{-1}]$ | $k_{34}$ $[s^{-1}]$ | $k_{43}[s^{-1}]$ | $k_{r1}$ $[s^{-1}]$ | $k_{r2}$ $[s^{-1}]$ | $k_{r3}$ $[s^{-1}]$ | $k_{r4}$ $[s^{-1}]$ | $\gamma$ $[s^{-1}][mol^{-1}]$ |
|---|---|---|---|---|---|---|---|---|---|---|---|---|
| *STL1* | $3.4 \times 10^{-3}$ | 5.1 | 1100 | $6.4 \times 10^{-3}$ | 1.5 | 8.3 | $3.7 \times 10^{-3}$ | $9 \times 10^{-5}$ | $2 \times 10^{-2}$ | 110 | $4.1 \times 10^{-2}$ | $5.3 \times 10^{-3}$ |
| *CTT1* | $3.8 \times 10^{-3}$ | 0.23 | 35 | $5.0 \times 10^{-3}$ | $4 \times 10^{-3}$ | $9.4 \times 10^{-3}$ | $4.4 \times 10^{-3}$ | $1.1 \times 10^{-3}$ | $9.9 \times 10^{-2}$ | $6.1 \times 10^{-1}$ | $2.4 \times 10^{-2}$ | $2.1 \times 10^{-3}$ |

TABLE II. HOG-Signaling Model Parameters

| $A$ − | $\alpha$ $[s^{-1}M^{-1}]$ | $\eta$ − | $m$ − | $r_1$ $[s^{-1}]$ | $\Delta$ |
|---|---|---|---|---|---|
| $9.3 \times 10^{9}$ | $1.45 \times 10^{-3}$ | 5.9 | $2.2 \times 10^{-2}$ | $6.1 \times 10^{-3}$ | $-5.31 \times 10^{-3}$ |

**SUPPLEMENTARY FIGURES**

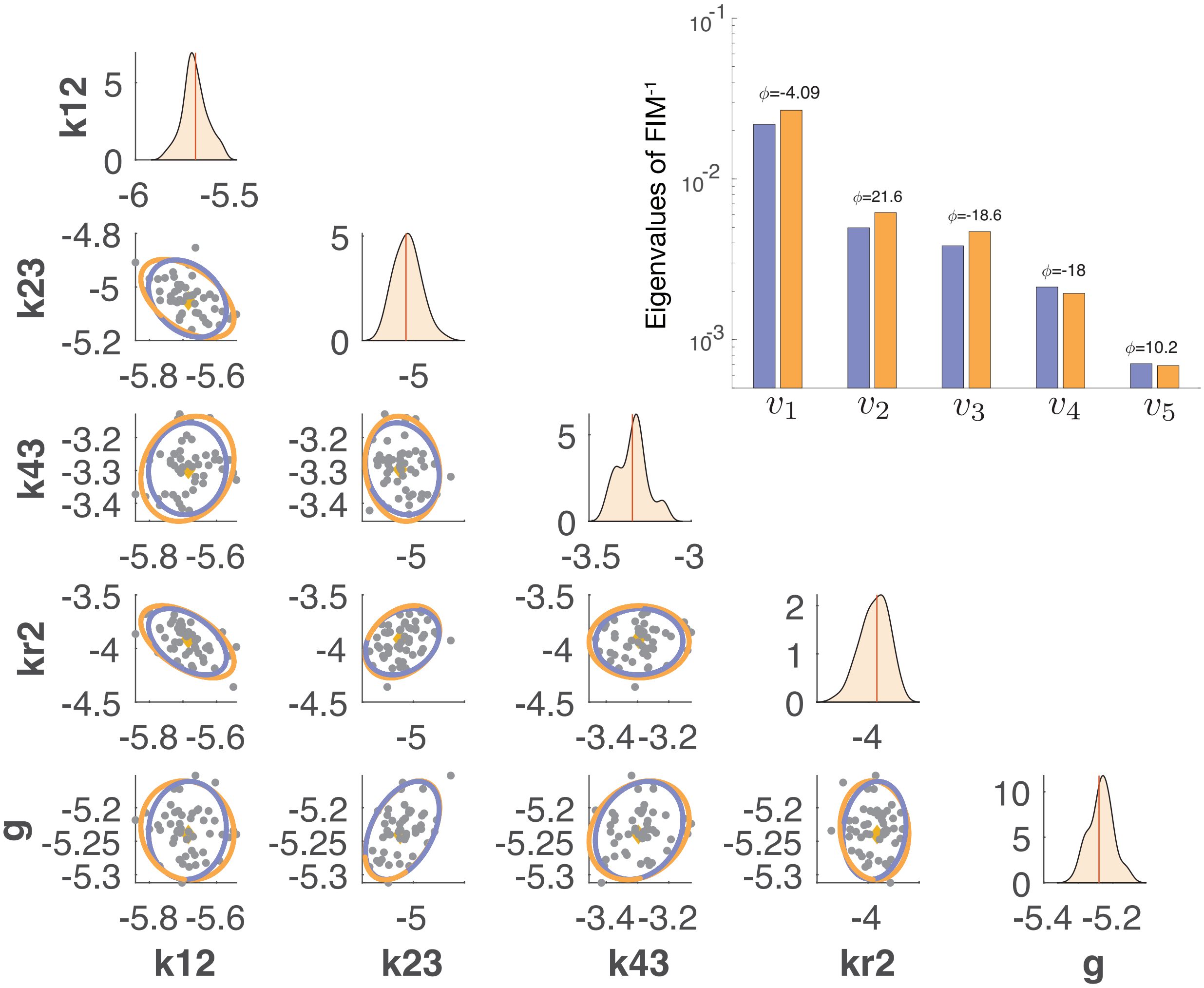


FIG. 1. *Verification of the FSP-FIM for the time-varying HOG-MAPK model.*(a) Scatter plots and density plots of the spread of MLE estimates for 50 simulated data sets for a subset of model parameters. All parameters are shown in logarithmic scale. The ellipses show the 95% CI for the inverse of the FIM (purple) and covariance of scatter plot (orange). The yellow dots indicate the parameters at which the FIM and simulated data sets were generated. (b) Rank-paired eigenvalues ($v_i$) for the covariance of MLE estimates (orange) and inverse of the FIM (blue). The angles between corresponding rank-paired eigenvectors ($\phi_i$) are shown in degrees.

[1] N. G. Van Kampen and N. Godfried, *Stochastic processes in physics and chemistry* (Elsevier, 1992).

[2] B. Munsky and M. Khammash, The Journal of Chemical Physics **124**, 044104 (2006).

[3] B. Munsky, Z. Fox, and G. Neuert, Methods (2015).

[11] B. Munsky, G. Li, Z. R. Fox, D. P. Shepherd, and G. Neuert, Proceedings of the National Academy of Sciences (2018).
